# A NEST-based framework unlocks massively parallel simulation of networks of multicompartment neurons with customizable subcellular dynamics

**DOI:** 10.1101/2025.06.30.662287

**Authors:** Willem A.M. Wybo, Leander Ewert, Charl Linssen, Pooja Babu, Elena Pastorelli, Pier Stanislao Paolucci, Abigail Morrison

## Abstract

While the implementation of learning and memory in the brain is governed in large part by subcellular mechanims in the dendrites of neurons, large-scale network simulations featuring such processes remain challenging to achieve. This can be attributed to a lack of appropriate software tools, as neuroscientific simulation software focuses on the one hand on highly detailed models, and on the other hand on massive networks featuring point-neurons. Here, we fill this gap by implementing a framework for the massively parallel simulation of simplified dendrite models with customizable subcellular dynamics. To achieve this, we leverage the NEural Simulation Tool (NEST), the neuroscientific reference with respect to efficient massively parallel simulations of point neuron networks. By co-opting the already existing model descriptions language NESTML, we generate C++ code implementing user-configurable subcellular dynamics. Through benchmarking and profiling, we show that the generated models run efficiently, leading to scalable NEST network simulations. We demonstrate relevant functionalities by showing that a key sensory computation – the association of top-down context arriving at distal dendrites in layer 1 and feedforward sensory input arriving perisomatically – can be achieved in a single shot fashion through apical calcium dynamics. Our work thus unlocks the study of how dendritic processes shape learning, and in particular of how brain-wide communication through long-range, layer 1-targetting connections steers perisomatic plasticity.

## Introduction

The brain computes in a massively parallel fashion. In the human brain, 86 billion neurons [1] continuously convert synaptic inputs into action potential (AP) output signals. Moreover, even at the subcellular level, computations proceed in a massively parallel fashion. Approximately 150 trillion synapses [2] (up to ∼ 10000 per neuron) are supported by complex molecular signalling networks within dendritic compartments. In itself, these signalling networks can also be understood as nanoscale computations that convert synaptic input, backpropagating APs, and local voltage and concentration signals into weight dynamics that support learning and memory [3]. Furthermore, the recent, explosive growth of AI technologies has shown the importance of scale [4], as it was achieved in large part by applying a well-known architectural motif [5] at massive scales [6–8]. Similarly, it can be expected that understanding the computational significance of subcellular processes will require embedding them in large-scale network simulations. It is only natural, thus, to use the parallelisation and vectorisation capabilities of modern supercomputing architectures to simulate the brain at scale in a massively parallel fashion.

Simulation of subcellular processes in biophysically detailed models has long been implemented in the successful NEURON simulator [9, 10], which is the most widely used tool for investigating models of individual neurons in great detail, and more recently in Arbor [11], which targets a similar use-case. Although NEURON admits efficient parallel simulations of network models through CoreNEURON [10], users have to take care of the parallel distribution of neurons across MPI processes themselves [12]. Moreover, NEURON’s API is geared towards complex models, providing interfaces that read morphological data files, or interfaces that construct neurons as combinations of cylinders [9]; the parameters of individual compartments are not exposed. Recently, however, there has been a rise in interest in capturing dendritic computational principles in a manner that is as simple as possible, to explore the role of dendritic computation at a systems level [13–18]. Therefore, an intermediate level of description is necessary, where parameters of individual electrical compartments are exposed, so that customized neuron layouts featuring few compartments and abstract dendritic spike mechanisms can be created.

For such purposes, the NEural Simulation Tool (NEST) is a promising candidate. It is the neuroscientific reference with regards to the massively parallel simulation of spiking network models [19], as the simulation kernel has been optimised to efficiently communicate spike times across parallel processes [20–23]. These capabilities generate little complexity for the user, as the distribution of neurons across MPI processes [24] and OpenMP threads [25] is taken care of by NEST itself. NEST has recently been extended with a GPU based framework for simulating spiking networks, further extending the capabilities for massive parallelism [23, 26, 27]. However, up until now NEST has had limited options to simulate more detailed compartmental models as parts of networks; users essentially had to develop custom C++ codes, or had to resort to using NESTML to explicitly define compartmental equations. The first option, however, is cumbersome, while the second option is suboptimal, as NESTML was traditionally aimed at single compartment models [28]. Standard NESTML will resort to an explicit Runge-Kutta integration scheme when compartmental dynamics are written explicitly [29], whereas a semi-implicit scheme based on the Hines algorithm [30] is more efficient and stable, and has been the gold standard for simulating spatially extended neuron models [9].

To facilitate the parallel simulation of subcellular processes embedded in neurons that are themselves embedded in large networks, we have leveraged the NESTML model description language [29] to develop a vectorized compartmental modeling framework in NEST. In this modelling framework, NESTML is used to describe dynamics that can, in principle, be simulated at each compartment of the neuron. In this sense, NESTML is thus used in a similar fashion to NEURON’s NMODL language [31,32]. These mechanisms can represent ion channels, synaptic receptors, concentrations dynamics, integrate-and-fire type mechanisms, and arbitrarily complex synaptic plasticity rules. The mechanisms can then be embedded at runtime into a user-configurable compartmental layout, which already provides passive dynamics by default. The structure of the tree, representing the dendrite and/or axon, is also configured at runtime. To simulate the model, the voltage at the next time step is computed by the Hines algorithm, while the user-defined mechanisms are simulated in a vectorized fashion. This means that each CPU can compute intracellular dynamics for up to eight compartments in parallel [33]. Under the hood, the standard NEST machinery takes care of distributing the neurons across threads or MPI processes, and of transmitting spikes between them, resulting in distributed network simulations of compartmental models that are efficient and easy to implement. In this paper, we demonstrate the excellent scaling performance of the resulting network models. We also provide a detailed description of the coding principles in the supporting information (SI), demonstrating that such massive network simulations can be achieved with little overhead to the user.

To illustrate the key principles, we implement a network model incorporating the observation that top-down afferents target distal dendrites in layer 1 and cause dendritic calcium (Ca^2+^) spikes [34–37]. We capture this computation in a simplified, two-compartment neuron that approximates the interplay between dendritic Ca^2+^ spikes and somatic action potentials (APs) [17]. We then demonstrate that this interplay is necessary and sufficient to achieve single-shot association learning between top-down context and feedforward sensory input when peri-somatic learning is governed by spike-timing dependent plasticity (STPD).

## Results

### Toolchain overview

Neuron models with complexity surpassing that of the point neuron are typically implemented using the multi-compartment (MC) modelling framework [38–41], where compartments are connected to each other through coupling terms:

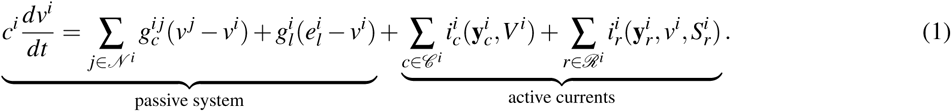

Here *v^i^* denotes the membrane potential in compartment *i*, *c^i^* its capacitance, *g_l_^i^* its leak conductance and *e_l_^i^* the leak reversal potential. The compartment *i* is coupled to its neighbours *N ^i^*through a coupling conductance *g^i^ ^j^*. Due to the conservation of current, the coupling is symmetric, *i.e.* 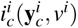. By identifying the compartments with the nodes of a graph and the neighbour couplings with the edges, a MC model representing a dendrite is always a tree graph.

An *a priori* arbitrary set *C ^i^* of ion channels may be present in each compartment *i*, as well as an *a priori* arbitrary set *R^i^* of synaptic receptors. Channel currents *i*_*c*_^*i*^(**y***_c_^i^*, *v*^*i*^) typically depend on the local voltage, and also on an *a priori* arbitrary set of state variables **y***_c_^i^*. These state variables may simply describe channel opening based on the voltage, such as in the Hodgkin-Huxley formalism [42], or may represent more complex intracellular dynamics [43–45], such as a dependence on certain ionic concentrations [46] or metabotropic interactions [47, 48]. Compared to ion channels, receptor currents 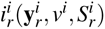 may similarly depend on complex intracellular dynamics through a set of state variables **y***_c_^i^* , to e.g. describe receptor dynamics and plasticity [49], but have the additional feature that they integrate incoming trains of action APs.

Leveraging NEST and NESTML, we have built a vectorized framework for the massively parallel simulation of spiking neural networks, where the neurons are multi-compartment models that can be equipped with arbitrarily complex dynamics. In this framework, active currents (cf. eq. (1); Fig 1A, left) are defined in NESTML, and compiled into vectorized C++ code (Fig 1B, left), allowing modern CPUs integrate currents for up to eight compartments in parallel [33]. The passive system (cf. eq. (1)) is assumed to always be present, and does not need to be described explicitly in NESTML. At runtime, after compiling the NESTML model, the dendrite structure is defined (Fig 1A, middle) while building the model, at the same time also setting the parameters of both the passive and active currents. Internally, this layout is translated to a tree graph data structures, which is also used to perform the Hines update of the model (see methods ; Fig 1B, middle)

**Figure 1.**
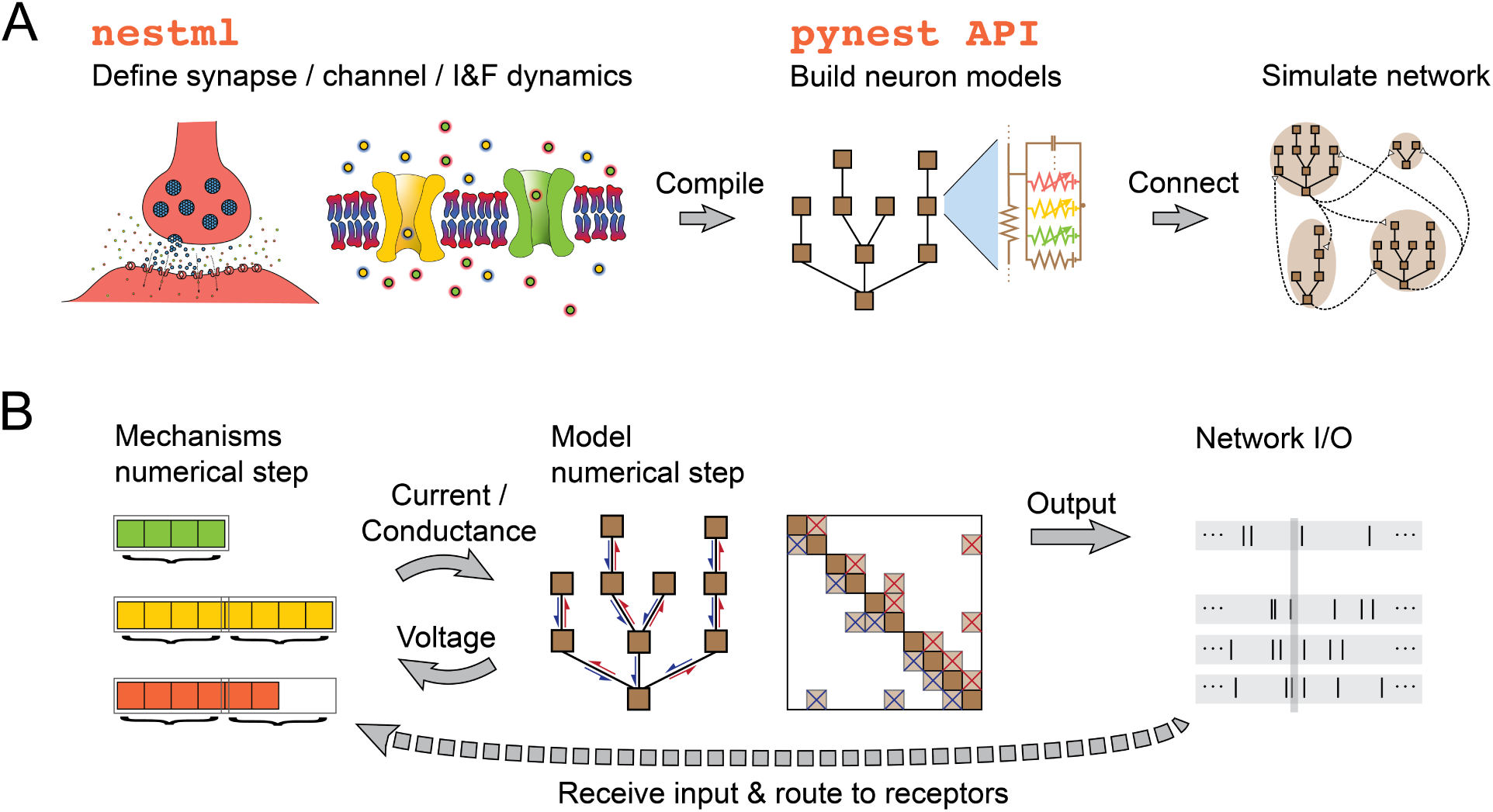
Overview of the compartmental simulation toolchain. **A:** Dynamics of transmembrane currents, such as voltage-dependent ion channels or synaptic receptor channels, are defined in the NESTML modelling language (left). These dynamical models are then compiled into efficient C++ code to simulate them as part of a compartmental model, as in eq. (1). The layout of this compartmental model is set in the PyNEST API (middle), including which channels and synapses are expressed in which compartments. Finally, this model is connected with other models to create a network simulation (right). **B:** The internal layout of the model consists of *(i)* a vectorized membrane current integrator for every current type defined in NESTML (left), *(ii)* a tree graph structure to advance the compartmental model through the Hines algorithm (middle) and *(iii)* a network API. The membrane current integrator reads out the voltage and relevant concentration variables from the compartments for which the current is defined, and uses the CPU vectorization to integrate the currents for multiple compartments in parallel. The results of the integration are then provided as the currents/conductances required for the implicit Euler integration of the compartmental model. External inputs are connected to the model through specification of a receptor ID, referring to the index in the receptor list, which is then mapped internally to the correct mechanism.

Finally, networks are created by connecting presynaptic neuron populations to specific receptor ports of the post synaptic neuron populations (Fig 1A, right). The incoming spikes are then routed internally by the model to the appropriate receptor current (Fig 1B, right).

### Single neuron and network scaling benchmarks

To test the efficiency of our generated code, we create two artificial morphologies where we can change the number of compartments at will. These morphologies also constitute two limit cases for the Hines algorithm: one of a fully branched model with one compartment per branch (Fig 2A, top) and one of a fully linear dendrite with all compartments on a single branch (Fig 2A, bottom). Note that for the linear dendrite, the morphology itself remains the same while the different compartment numbers represent a different spatial discretization. For the branched model, the number of dendritic branches changes with the number of compartments, resulting in a higher effective leak and capacitance. While these effects do influence input integration, they are only of relevance for the network benchmarks, as the number of transmitted spikes may affect network runtimes but – in case of Hodgkin-Huxley type channels – does not influence single neuron runtimes. We equip all of the compartments with a sodium and potassium channel, and defined identical models in NEURON and NEST, as can be seen by a comparison of their membrane potentials (Fig 2B). We then simulate these models for increasing numbers of compartments, and measure the runtime of a simulation of 10s (Fig 2C) on 2.25 GHz AMD EPYC processors that are part of the JURECA DC module [50]. The simulation time of NEST is less than that of NEURON for all tested numbers of compartments with a greater disparity at greater numbers: the speed-up of NEST with respect to NEURON increases from a factor of two for single compartment models to a factor of four to five for models with 100 compartments or more (Fig 2D). To understand how the runtime of the model is distributed over the various computational steps, we profile the linear dendrite model (Fig S1A), which reveals that solving the Hines algorithm takes up only a minor fraction of the runtime (up to 17%, Fig S1B), whereas most of the time is spent updating the active mechanisms, and specifically computing the exponentials necessary for that purpose.

**Figure 2.**
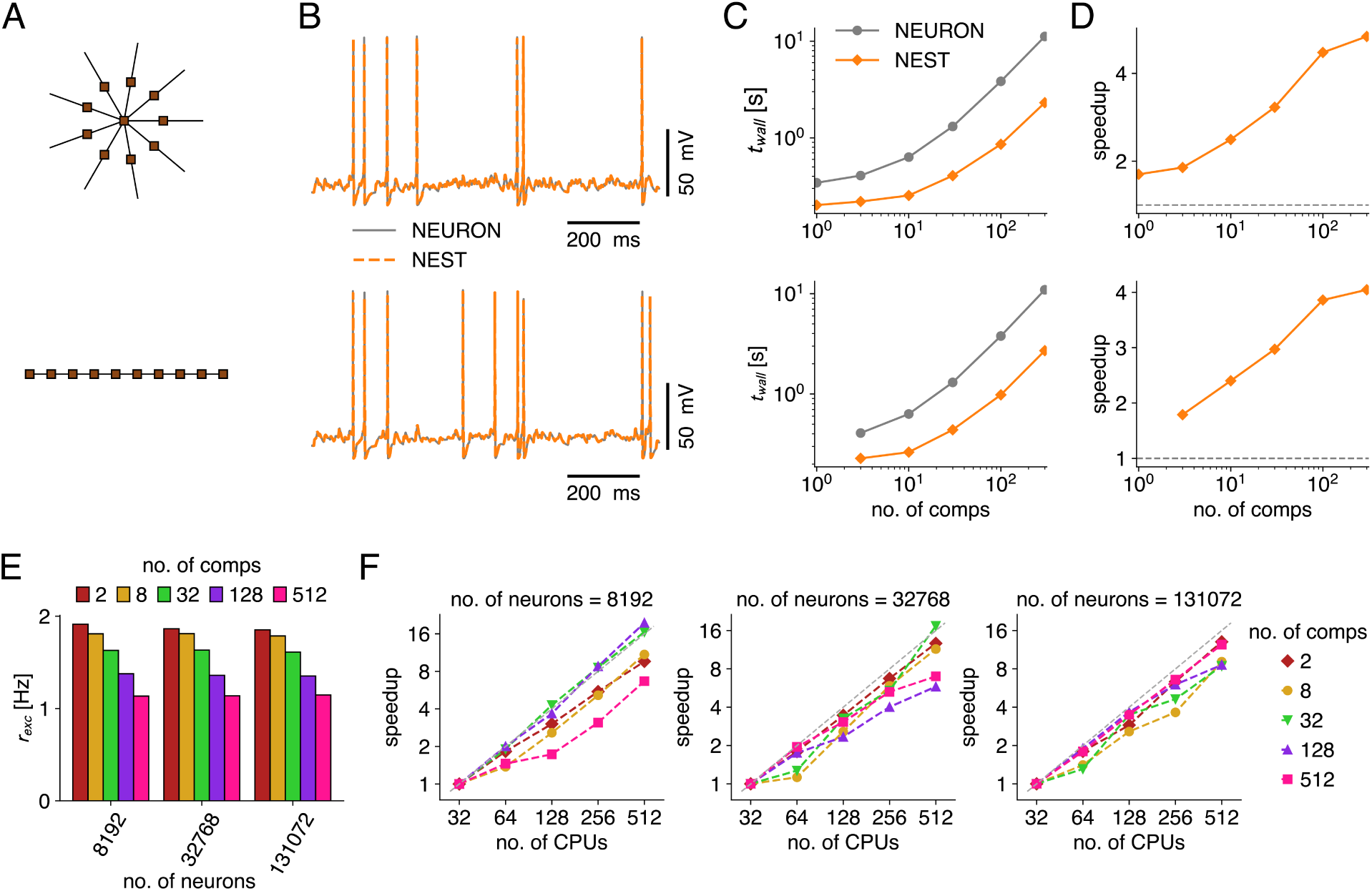
Benchmarking of the NEST-based compartmental modelling framework. Comparison of single neuron wall clock times between NEST and NEURON (A-D) and assesment of network scaling in NEST (E,F). **A:** Two constructed morphologies allow precise control of compartment numbers and represent edge cases of the Hines algorithm: a star layout where each dendrite consists of a single compartment (top) and a linear layout without branching (bottom). These morphologies are equipped with Na- and K-channels and simulated in both NEST and NEURON. **B:** Comparison of somatic voltage traces between NEST and NEURON for the star morphology (top) and linear morphology (bottom), which both contain ten compartments in these simulations. **C:** Wall clock time comparison between NEST and NEURON for a simulation time of 10 s as a function of the number of compartments for the star (top) and linear (bottom) morphology. **D:** Speed-up factor of NEST with respect to NEURON. **E:** Firing rate of the network configurations. In these configurations, varying numbers of neurons with different compartment numbers (using the star morphology, A, top) are recurrently connected with fixed in-degree, and stimulated with Poisson processes of fixed rate, resulting in comparable firing rates across configurations. **F:** Strong scaling performance of the network models, i.e. speed-up measured in wall clock runtime when the same network is distributed over increasing amounts of CPUs, for a fixed simulation time of 10 s. Panels show and increasing number of neurons (left 2^13^, middle 2^15^, and right 2^17^), and in each panel different compartment numbers per neuron are compared (8, 32, 128, and 512). The dashed grey line shows the theoretical optimum, where increasing the number of CPUs by a given factor results in an equal speed-up.

We furthermore test the scaling performance of a recurrently connected spiking network, in which the neurons have the star morphology (Fig 2A, top). Networks are simulated with increasing numbers of neurons (2^13^, 2^15^, 2^17^), and for each network size we probe four different numbers of compartments (8, 32, 128, and 512). Approximately 20% of neurons are inhibitory, while the other 80% are excitatory. To minimize network size effects, we randomly connect these neurons with a fixed in-degree, and also connect Poisson spike generators to the excitatory population, likewise with a fixed in-degree. Note that the dendritic compartment receiving the connection is also chosen at random. Furthermore, we apply a weight normalization based on the input resistance, to limit the effect of neuron size induced by varying the number of compartments in the star layout on output firing rate. This results in network states with firing rates that were independent of the number of neurons, but still changed somewhat between ∼ 1 to ∼ 2 spikes/s for the different numbers of compartments (Fig 2E). We consider these firing rate variations sufficiently minor, so that runtimes of the full simulation would only marginally be affected by spike transmission. We then measure the wall clock runtimes of network simulations of 10 s, while distributing them over increasing numbers of CPUs (Fig 2F). We found that runtimes increased approximately linearly with numbers of neurons as well as with numbers of compartments (Fig S2), as expected. Furthermore, the speed-up as a function of the number of CPUs for fixed network and neuron size is close to the theoretical optimum for strong scaling experiments (Fig 2F). This excellent scaling performance was observed for all network sizes, with as few as 16 neurons per CPU, indicating that large-scale networks can be distributed over many CPUs with little overhead. Furthermore, achieving these distributed network simulations in NEST is straightforward and requires only few lines of code, as NEST natively takes care of distributing the simulation across MPI processes and/or openMP threads.

### Hybrid few-compartment models uniting detailed and abstract dynamics

A major use-case for our simulation framework is defining hybrid MC models that combine abstract integrate-and-fire dynamics representing AP and/or dendritic spike generation with select biophysical processes of interest (Fig 3A). Through the NESTML description language [29], the equations governing the dynamics of these models can be defined compactly (Fig 3B), while a scalable network simulation can then be achieved easily with NEST, as argued in the previous section. We illustrate these principles in a model uniting adaptive exponential (AdEx) integrate-and-fire dynamics [51] and dendritic calcium dynamics (Ca-AdEx), which we base on our prior work [17]. These calcium dynamics, which drive Ca^2+^-spikes, couple top-down inputs in the distal apical dendrites of pyramidal neurons to the feedforward inputs impinging peri-somatically [34–36, 52] (Fig 3A, left). The resulting Ca^2+^-spikes are implicated in computations ranging from error backpropagation to context adaptation [53–57], and require a calcium-mediated interplay between calcium channels and calcium-activated potassium channels, next to backpropagating APs supported by dendritic sodium channels [58]. In this hybrid, two-compartment description, we can not support backpropagating APs in the same way that biophysically detailed models do, where the apical dendrite is modeled through a multitude of compartments. Instead, backpropagation of APs to the dendritic compartment is incorporated explicitly (Fig 3B, lines 13-15). We test this model by reproducing the backpropagation-activated coincidence detection protocol [34] (BAC-firing), where the model spikes once in response to a somatic stimulus, does not spike in response to a dendritic stimulus, but fires a burst of APs in response to the coincidence of both (Fig 3C).

**Figure 3.**
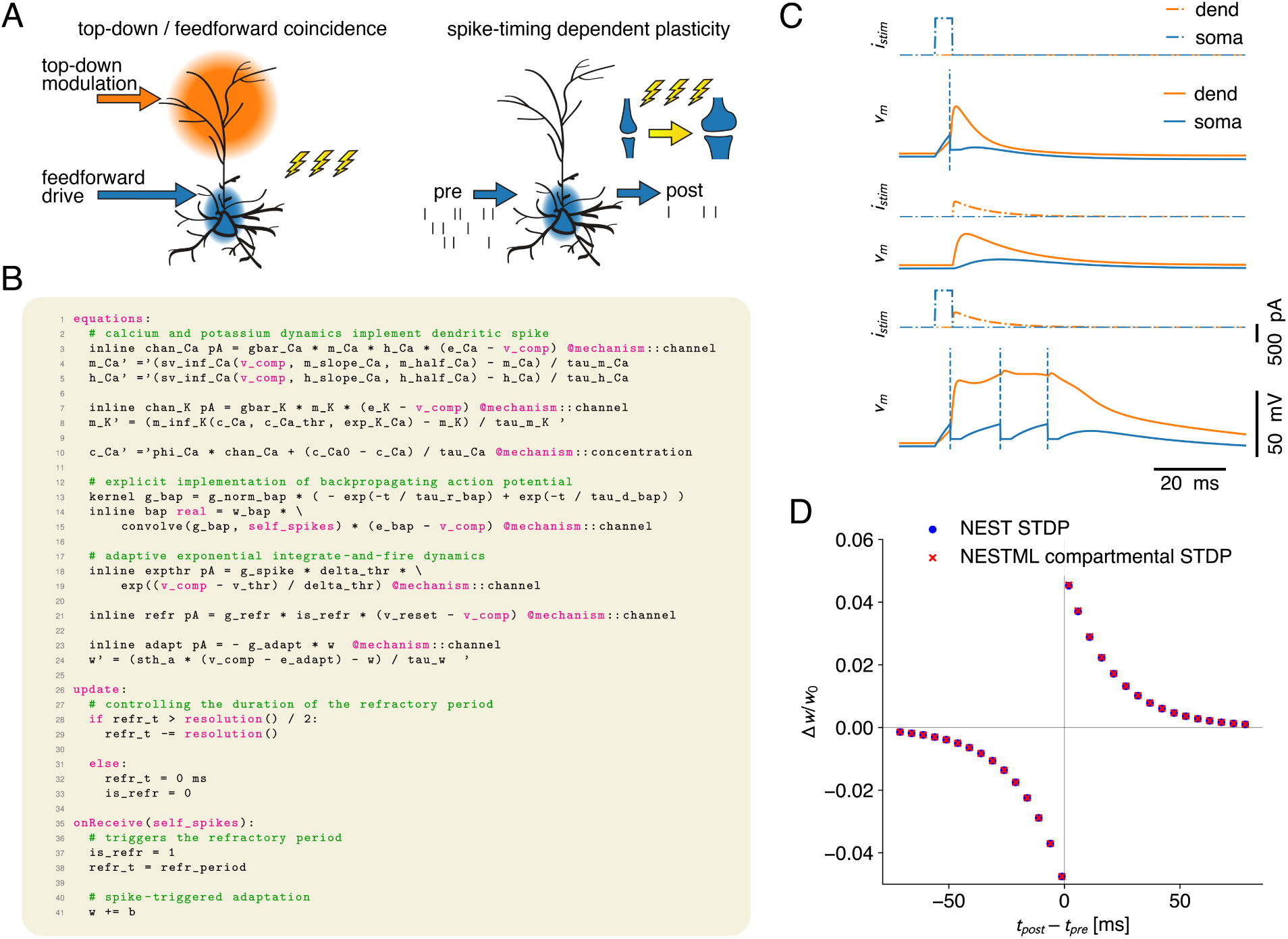
Exemplar implementation of a hybrid model uniting dendritic calcium dynamics and a somatic adaptive exponential integrate-and-fire mechanism. **A:** Coincidence of apical top-down input and peri-somatic feedforward input elicits BAC-firing, a calcium-mediated coincidence detection mechanism (left). In a two-compartment NESTML model, this is combined with somatic STDP (right). **B:** NESTML code excerpt of the Ca-AdEx model, which features the Ca^2+^-dynamics necessary for BAC firing, as well as AdEx dynamics for somatic spike generation. Note that the NESTML blocks specifying parameters and state-variables, as well as the NESTML model implementing STDP, are omitted for clarity. SI and Figs S3-S6 provide a more complete overview of the essential code concepts. **C:** Reproduction of the BAC-firing protocol with the two-compartment Ca-AdEx model, where the model spikes once in response to a somatic stimulus (top), does not spike in response to a dendritic stimulus (middle), but fires a burst of APs in response to the coincidence of both (bottom). Somatic and dendritic input currents (blue resp. orange) are shown above the somatic and dendritic response traces (blue resp. orange). **D:** Reproduction of the STDP window with the two-compartment Ca-AdEx model, once using the standard NEST STDP mechanism (blue dot) and once using the NESTML one embedded in the MC framework (red cross).

While static weights permit the study of dynamical network states, understanding how biological brains learn requires the investigation of how neural activity and neurotransmitter signals drive weight changes. NEST provides a wide variety of plasticity rules for this purpose, which range from classic STDP [59, 60] to credit assignment rules for error-driven learning. These plasticity rules can straightforwardly be combined with the MC modeling framework (Fig 3A, right, Fig S6), where several options are available to the user. For instance, NEST’s built-in plasticity rules can be combined with any NEST model, including the compartmental ones. Alternatively, the MC framework also facilitates the definition of custom plasticity rules in NESTML, which can read out any compartmental state variables. We demonstrate this functionality by re-implementing the STDP rule in the MC framework, and combining it with the Ca-AdEx model (see SI), which results in equivalent dynamics to the standard NEST STDP implementation (Fig 3D).

### Leveraging dendritic dynamics for single-shot learning

While a broad consensus is emerging that dendritic dynamics fundamentally shape brain functions involving learning and memory [56, 57, 61–66], large-scale network studies of this are rare because of a lack of appropriate simulation tools. Our NESTML compartmental modelling framework allows us to straightforwardly implement network simulations featuring dendritic dynamics, which we demonstrate by embedding the previously described neuron reproducing dendritic Ca^2+^-spikes in a network model. One striking aspect of the cortical layout, is that contextual afferents from higher cortical areas target layer 1 [67], where they connect to distal apical dendrites of pyramidal cells and disinhibotory vasoactive intestinal peptide-expressing (VIP) interneurons [68]. Together, this allows for the generation of apical Ca^2+^-spikes, which fundamentally shape perception [37, 69].

Here we demonstrate, in a recurrently connected network consisting of the Ca-AdEx neurons, that contextual input impinging on the apical dendrite can drive the formation of experience specific neural assemblies in a single-shot fashion, following an apical stimulus lasting only 25 ms. Thus, in this set-up, apical input enables learning of individual experiences at a rate compatible with the fastest conscious biological perceptions [70, 71]. We construct a network model with four neural assemblies, where inhibitory interneurons implement a winner-take-all dynamic (Fig 4A).

**Figure 4.**
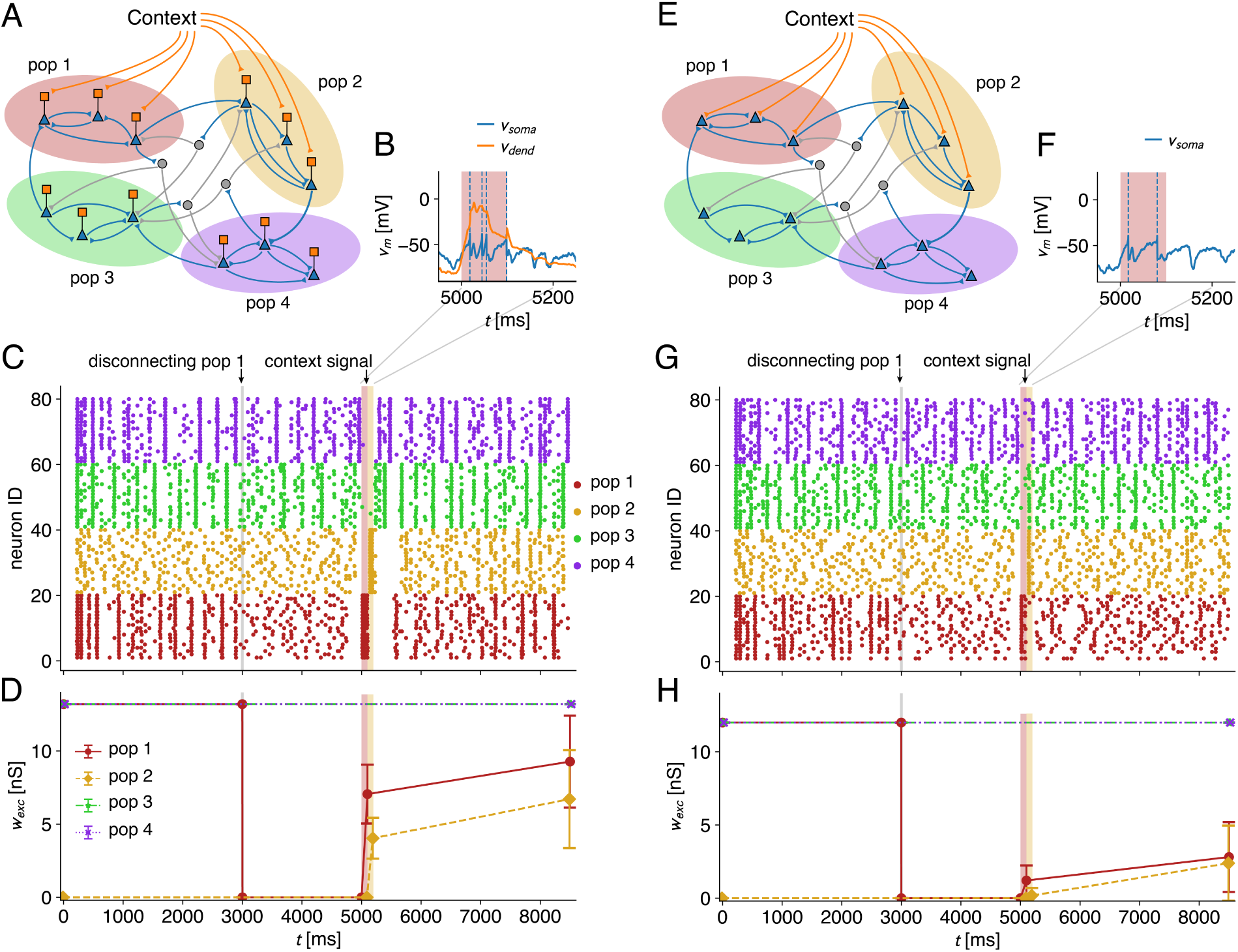
Dendritic, but not somatic contextual input drives single-shot association learning to yield a winner-take-all network model. **A:** A network with four populations of Ca-AdEx neurons is connected recurently through somatic connections, while inhibitory interneurons implement broad inhibition that results, for strong intra-population excitatory weights, in a winner-take-all network where each population fires in unison and inhibits all others. Contextual connections impinge on the dendritic compartment. **B:** The contextual input triggers a dendritic Ca^2+^-spike, resulting in somatic burst firing. **C:** Raster plot of the spiking network activity. Initially, populations 1, 3, and 4 have high intra-population connection weights, and therefore each firing in a concerted manner, while population 2 has zero connection weight and fires in and uncoordinated fashion. To illustrate the effect more clearly, connection weight in population 1 is set to zero at *t* = 3000 ms, resulting in random firing. A 25 ms context signal, in the form of Poisson input, is then delivered first to population 1 at *t* = 5000 ms, and then to population 2 at *t* = 5100 ms, while STDP is turned on in the intra-population connections of populations 1 and 2 at *t* = 5000 ms. **D:** Corresponding intra-population weights in the network. **E-H:** Same as A-D, but for point neurons, where the context signal has to target the somatic compartment.

With recurrent connections impinging perisomatically, neurons belonging to an assembly fire in unison, inhibiting all other assemblies (Fig 4C). Conversely, weak intra-cluster connections lead to random firing, as is exemplified by setting the intra-assembly connection weights for population 1 to zero at *t* = 3000 s (Fig 4D). We then target the the apical compartments in the randomly firing clusters (populations 1 and 2) with excitatory contextual inputs. To mimick VIP-mediated disinhibition, we provide inhibitory inputs to the apical compartment in the 200 ms preceding the excitatory input, so that the release from inhibition occurs when the excitatory input arrives. Thogether, this elicits a Ca^2+^-spike in the targeted populations (Fig 4B), in turn triggering burst firing in the associated neurons. In combination with STDP plasticity on the intra-assembly connections, this burst leads to rapid potentiation of these synapses (Fig 4D), and restores coordinated ensemble firing (Fig 4C). When the network consists of point neurons (Fig 4E), the same context signal as before has to impinge perisomatically, and hence no burst firing is elicited (Fig 4F). As a consequence, the context signal does not drive potentiation and leads to a failure to obtain ensemble formation in a single shot fashion (Fig 4G, H). Therefore, in neurons missing the apical amplification mechanisms, either a contextual stimulation of much longer duration or of unrealistically strength would be required for single shot learning at rates of ten frames per second. In many studies so far, only such point-neuron implementations were investigated, as only these were available for straightforward large-scale simulation. Our results therefore also highlight that mechanistic conclusions about brain computation obtained from point-neuron network simulations should be treated with care.

## Discussion

In this work, we have presented a novel framework for creating large-scale network simulations consisting of multi-compartment neuron models, leveraging the spiking neural network simulator NEST [72]. This framework is based on the existing model description language NESTML [28, 29, 73, 74], exploiting its syntax features to describe membrane currents and concentration dynamics. We have also extended NESTML with a new code generation pathway that builds efficient multi-compartment C++ code based on the provided equations. The models can then be configured in the PyNEST API and connected in networks to be simulated using the standard NEST routines. Through the use of a structure-of-arrays layout, the models are efficiently vectorized, leading to considerable single-neuron speedups with respect to the NEURON simulator. Together with the native MPI support for massively parallel network simulations built into NEST, this results in a framework that allows the specification and simulation of networks with dendritic dynamics at scale.

Although there are no a-priori restrictions on the amount of detail and the number of compartments a model can have, the model configuration interface in the PyNEST API has deliberately been kept low-level, i.e. the parameters for the individual compartments have to be explicitly provided by the user. This interface is hence well-suited to multi-compartment models containing few compartments, which are increasingly used to capture key dendritic computations in a minimal fashion [13–18]. However, the NEST initiative has also recently been extended with the NEural Analysis Toolkit (NEAT) [75], a novel tool that allows the researcher to define morphologically detailed neuron models, analyze them, and create simplifications thereof. These fully-detailed and simplified neuron models can then be exported to NEURON and NEST. To achieve the latter, NEAT leverages the same NESTML based code-generation pipeline and C++ framework as described here, resulting in the efficient simulation of networks of biophysically detailed neuron models.

By enabling fine-grained parallelization and vectorization, GPUs are highly adept at implementing computations requiring many identical operations [76]. This renders them ideally suited for e.g. matrix multiplication, leading to large speed-ups for simulating artificial neural networks compared to normal CPUs. For spiking neural networks, the situation is more nuanced, as speed-ups in this case have been reported to be on the order of 1.5 to 3 [23, 26, 27], by comparing NEST GPU and GeNN [77, 78] to normal NEST. Together with the wide available of multi-CPU systems, this presents massively parallel CPU-driven simulations of spiking networks as a competitive and cost-effective approach. Nevertheless, these performance results were obtained for single compartment models. Likely, the GPU speed-up with multi-compartment models will increase with increasing compartment numbers [79]. For this reason, the NEST initiative intends to also provide code generation for the multi-compartment models in NEST GPU, as is already being developed for point neuron models.

Converging evidence [34–36] links the cellular and architectural organization of the cortex and hippocampus to neural dynamics across wake and sleep. At the cellular level, the distinct morphology of layer 5 cortical pyramidal neurons, with separate apical tuft and peri-somatic zones, enables compartment-specific integration of inputs. On a larger scale, inter-areal projections preferentially target the apical tufts, conveying contextual or internal signals, while sensory inputs predominantly impinge on the (peri-)somatic region. This organization points to apical amplification [80–82], where during wakefulness selected neurons are boosted into high-frequency bursting in response to coincidences between top-down priors and bottom-up input. In turn, this amplification supports the formation of sparse, experience-specific assemblies at a faster rate than would be possible in point-neurons. In contrast, during deep sleep, apical isolation is thought to decouple these dendritic compartments, reducing integration and enabling the optimization of local circuits [83]. The efficacy of learning algorithms based on apical mechanisms has been demonstrated in networks either by using abstractions of multi-compartment neurons [57, 84, 85], or by emulations in models composed of single-compartment neurons [86,87]. These appraoches however do not generalize to large-scale spiking networks due to the missing calibration of contextual and perceptual currents at growing scales, which is enabled by true simulation of apical mechanisms. The present work unlocks the study of incremental learning based on this apical mechanism in large-scale spiking networks, which in the long run could lead to improved artificial intelligence systems leveraging this biological mechanism.

Recent years have seen a surge in the capability of artificial intelligence systems [6–8]. This can largely be attributed to an increase in scale [4], both in terms of network size and compute, as the base network architectures have not fundamentally changed [5]. To elucidate their role in shaping high-level computations, dendritic trees will similarly have to be embedded in large scale simulations implementing learning and memory. Only then can their computational role at the network level emerge. The work presented here implements the simulation technology necessary for this endeavour, by proposing a flexible framework that enables the simulation of large-scale networks with dendrites in a massively parallel fashion. While the current implementation focuses on CPU systems with NEST, the use of the NESTML framework will enable code generation for GPUs as well as for a variety of emerging neuromorphic computing systems [88, 89].

## Acknowledgements

This work was supported by Helmholtz Association’s project-oriented funding programme (PoF 2, Topic 3). Also, it has been co-funded by the European Next Generation EU through Italian MUR grants CUP I53C22001400006 (FAIR PE0000013 PNRR Project) and CUP B51E22000150006 (EBRAINS-Italy IR00011 PNRR Project). The authors gratefully acknowledge the computing time granted on the supercomputer JURECA [50] at Forschungszentrum Jülich under grant jinm60. Further, we would like to extend our gratitude towards Christophe Blaszyck and Jakob Jordan for contributions to the initial prototypes.

## Code Availability

Code to reproduce the simulations in this paper is available at https://github.com/WillemWybo/WTA-states/ releases/tag/v1.0 for the winner-take-all network and at https://jugit.fz-juelich.de/w.wybo/ca-adex for all other simulations.

## Footnotes

1 https://jinja.palletsprojects.com/en/stable/

## Methods

### Numerical integration of the compartmental dynamics

The membrane voltage in a multi-compartment neuron model, i.e. a model that follows (1), is integrated using the field-standard implicit Euler approach [9], [90], which ensures that the integration is stable. To that purpose, eq. (1) is discretized in the following way:

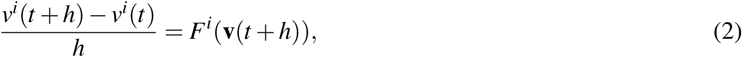

where *h* is the integration step and *F^i^* the entire right-hand side of eq. (1) divided by the capacitance *c^i^*. Here, the unknown voltage **v**(*t* + *h*) = *v*^1^(*t* + *h*)*,…, v^N^*(*t* + *h*) at time *t* + *h* is featured inside the complicated nonlinear function **F**(**v**) = *F*^1^(**v**)*,…, F^N^*(**v**). We linearize this function around the known voltage **v**(*t*), i.e.

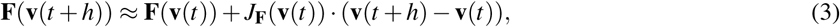

where *J***_F_** is the Jacobian of **F**. By substituting (3) in (2), it follows that solving the matrix equation:

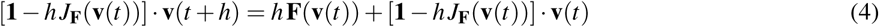

yields the next timesteps’ voltage **v**(*t* + *h*). To efficiently solve this equation, the generated C++ code leverages the Hines algorithm [30] (Fig 1B, middle). For the state variables of ionic channels and receptor currents, we employ the leap-frog scheme: a state variable *y* is computed at 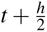 , and thus has this value in both *F^i^*(**v**(*t*)) and *F^i^*(**v**(*t* + *h*)). Conversely, to compute the time evolution of a state variables from 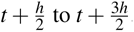 , the voltage *v^i^*(*t* + *h*) is taken to be constant over this time-step. The standard NESTML pipeline, featuring the ODE-toolbox [28], is used to determine the best method of advancing the state variables. In many cases, this results in analytical integration to the next time-step (e.g. typical for Hodgkin-Huxley type channels) using a method known as exponential Euler [91]. In more complex cases, the C++ code implements the 5th order Runge-Kutta scheme [90].

### Code generation pathway

The NESTML code generation pipeline follows the standard NESTML pattern [29] into which we inject our additional processing at specific steps. This general pattern starts with the initialization, where the correct code generator and other settings are configured based on user input. After that the model is parsed, syntactic and semantic checks are performed. Then the actual code generator is invoked which inherits from a base NESTML code generator class.

**Model parsing.** The purpose of parsing the model is to verify the code’s syntactic correctness and to translate the NESTML code into an Abstract Syntax Tree (AST), a data structure that can be easily manipulated and traversed. The AST organises all instances of syntactic elements that appear in the code into a hierarchical structure, and comes with a `ASTVisitor` class that, when called upon, traverses the tree recursively from any given node, using the “visitor” design pattern. For each `ASTNode` type, one can define visitor functions which can modify node content and extract information.

**Mechanism-specific information collection.** In the normal NESTML code generation context conditions checks (CoCos) verify that the NESTML code is semantically valid after building the AST. However, the compartmental case requires further information collection, as we need to distinguish between the various types of mechanisms. The original AST structure is not helpful to store this information, because it does not organize elements by their mechanism ownership. Since certain semantic checks can only be done when the ownership of elements is revealed, we inject our information collection into the CoCo process.

Each type of mechanism has its own information collector, which inherits from a base collector. From the inline that defines a mechanism – i.e. the ones marked with the @mechanism::{type} tag (cf. SI & Fig S3) – we recursively search all referenced variables, their initialisation expressions, functions, equations and kernels. From the NESTML update block and our special case of the `onReceive()` function for self-spiking, we extract the parts associated to the mechanism. Finally, we utilize ODE-toolbox to find solutions for the ODEs referenced per mechanism. This general collection process is applied to each mechanism. For the channel and concentration mechanisms, we furthermore check whether there are parameter combinations which result in inputs that are identically zero, so that we can omit their simulation from those compartments. This is a key aspect of the modularity of the compartmental feature.

The synaptic processing performs very similar information collection as is the case for neurons in terms of the structure of collected elements, states, functions, ODEs and so on. We don’t insert this into the CoCos but instead into the original synapse processing because we don’t perform additional semantic checks that rely on a mechnism-wise distinction of elements as is the case for neurons. In addition, we collect the update block, the spiking ports, determine which is the pre- and post-synaptic port based on the code generator options, and collect the `onReceive()` blocks for those ports. We collect variables that potentially refer to a postsynaptic neuron variable. As for the neurons, we utilize part of the original transformations for inlines and convolutions.

**Converting equations to C++ expressions.** After the CoCos have been performed successfully, the code generator is invoked for each mechanism that has been parsed. Here, we start by invoking the parts of the standard neuron processing pipeline where the incoming spike buffers for synapses are generated, inline references are replaced with their expressions, and convolution calls are replaced by a new variable that will contain their result.

We insert an additional step into the template namespace setup. The namespace is a dictionary that contains all the data and Python functions referenced inside the template. In this additional step, we perform modifications to AST nodes in our collected mechanism dictionaries, which we could not perform during the original collection, as it requires the invocation of the NESTML parser, which in turn invokes the CoCos, leading to a circular dependency.

During this information enrichment process, we parse the resulting expressions from ODE-toolbox, add the propagators to the internals block, and collect the updated expressions in which the inlines and convolutions have been substituted. We need to parse our new expressions and all new resulting variables as AST nodes because the already defined C++ printers in NESTML rely on this input format.

### The C++ code structure

We generate the C++ code with Jinja1, a templating language that can process standard Python data structures and to which one can pass Python objects that can be used in the Jinja logic blocks. Jinja also provides control structures with a Python-like syntax.

**The generated C++ files.** A module, as defined by the user, may include an arbitrary number of models that can be passed to the generator. The class that registers the module is an extension of the `NESTExtensionInterface` class in which the contained models are registered with the simulator. The name of this generated file is `<MODULE_NAME>_module.cp`

The top model classes are defined in `<MODEL_NAME>_nestml.cpp/.h` These inherit from the NEST `ArchivingNode` class. When writing a custom NEST model in C++, one must define certain functions in the model class like `update()`, `pre_run_hook()` and `set_status()`. The `update()` function is called at every timestep during the simulation. The `pre_run_hook()` function may be used to internally configure the neuron properly before the simulation starts. The `set_status()` function may be called explicitly from the Python interface or is called implicitly when assigning a member variable of the Python neuron object. It receives a dictionary with entries corresponding to the user-defined or certain predefined states and parameters of the neuron. In our case we also use this function to add new compartments and receptors to the compartmental tree by detecting an entry with id “compartments” or “receptors”, which internally is a dictionary in which we expect to find the parent node id and a corresponding parameter dictionary.

In the `cm_tree_<MODEL_NAME>_nestml.cpp/.h` files the compartmental tree data structure is defined. This class represents the user-defined tree structure of the model through `CompartmentNode` attributes that contain all parameters and states that are part of the passive dynamics, which are implicitly included in all compartmental models. The tree class as container owns the root compartment and has a list of pointers to all compartments in the order in which they were added. This class integrates the passive dynamics directly and triggers the integration of compartmental dynamics not through the compartment class but through the `NeuronCurrents` class. Finally, it combines the integrated compartment internal dynamics and performs the integration of cable dynamics through the Hines algorithm.

The `NeuronCurrents` class is defined in the `cm_neuroncurrents_<MODEL_NAME>_nestml.h` file. It primarily contains an update function that combines integration results from all mechanisms across the neuron and manages the instances of mechanisms that the user adds to the compartments of the neuron. The `cm_neuroncurrents_<model_na` and `.cpp` also define the mechanism classes, which are generated based on the mechanisms that have been defined by the user. Each mechanism class defines three functions that are relevant to the core functionality. The `f_numstep()` function, which is called at every update of the simulator, integrates the defined dynamics. Synaptic dynamics are added to the `f_numstep()` in case of the receptor-synapse mechanisms. The `pre_run_hook()` function, precomputes quantities that are not dependent on any evolving state. And a `f_self_spike()` function, which is the reduced form of the `OnReceive(self_spike)` function in the model, which is always called when the neuron spikes. In all mechanism types, except for the concentration type, the `f_numstep` function returns the current induced by the mechanism.

**Micro-parallelization through vectorized C++ code.** One of our goals was to achieve a further level of parallelisation – beyond the MPI OpenMP parallelisation that NEST implements at the network level – by vectorizing the integration of mechanism dynamics. Vectorization is a parallelisation technique that relies on a so-called vector register on the CPU, which can contain multiple values of the same type on which the same operation is performed in one step. To make this possible, the data on which computation is performed needs to be provided in the form of an array or array container, such as the vector datatype in C++. We vectorize all instances of the same mechanism type added to the whole neuron. Each object of a mechanism class thus encapsulates all its instances for the entire neuron, except for those cases in which the parameters result in a contribution that is identically zero (up to machine precision). Each defined state and parameter is stored as a vector, where each entry contains the state or parameter value in a single compartment. When integrating the associated dynamics, we can loop through all instances. By marking this loop with `#pragma omp simd`, we tell the compiler to vectorize the operations contained in this loop. Additionally, we need to consider that we also call functions within this loop. These functions need to be vectorizable. We found that we need to do the following to get the compiler to consistently vectorise our functions. When declaring these functions we need to add the pragma `#pragma omp declare simd` and mark them as inline with `attribute` `((always_inline))` in addition to the `inline`keyword. Furthermore, we need to compile with the flags `-march=native` and `-ffast-math`. We are aware that the `-ffast-math` flag may introduce numerical artefacts due to the nature of floating-point arithmetic, but in our tests, we did not encounter any issues.

### Simulation details

**Benchmarking and scaling experiments.** All benchmarking and scaling experiments w ere performed on the Jureca supercomputer at Forschungszentrum Jülich [50], except for the profiling experiments, where we used a laptop with with Intel(R) Core(TM) i5-4210H CPU @ 2.90GHz and 16 GB DDR3 1600MHz Ram. In the comparisons with NEURON, we used NEURON version 8.2.6.

The artificial morphologies have a soma with radius 10 *µ*m. The star neuron, for *N* compartments, was then equipped with *N* − 1 cylindrical branches of length 10 *µ*m and radius 1 *µ*m, which each were converted into a single electrical compartment using standard geometrical principles [9]. The linear neuron had a single dendritic branch of radius 1 *µ*m and length 1000 *µ*m, which was discretized in *N* − 1 compartments. Na [92] resp. K [93] currents where distributed at the soma with densities 1.71 · 10^6^ *µ*S/cm^2^ resp. 0.766 · 10^6^ *µ*S/cm^2^, and with 1/10’th of the somatic density in the dendrites. Finally, passive leak conductance and reversal were fitted to obtain a membrane time-scale of 10 ms at an equilibrium potential of 75 mV.

Synaptic conductances in most simulations followed the dual exponential profile [94]:

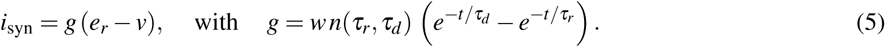

with parameters *τ_r_*= 0.2 ms, *τ_d_* = 3 ms, and *e_r_*= 0 mV they represented excitatory (exc) AMPA synapses, whereas with parameters *τ_r_* = 0.2 ms, *τ_d_* = 5 ms, and *e_r_* = −80 mV they represented inhibitory (inh) GABA synapses. *n*(*τ_r_, τ_d_*) is a normalization factor so that the maximum of the conductance windows is the synaptic weights *w*.

In the single-neuron benchmarks, we only used AMPA synapses, one in each compartment, and adjusted the input Poisson spike rate such that the total input rate to the neuron was 200 Hz. Excitatory weight *w* was 4 nS. In the network benchmarks, we also used inhibitory neurons and thug GABA synapses, and had five connection weights: *w*_exc→exc_ = 1 nS, *w*_exc→inh_ = 2 nS, *w*_inh→exc_ = 8 nS, *w*_inh→inh_ = 2 nS, and *w*_ext→exc_ = 2 nS. The neurons in the network where connected randomly with fixed in-degree. The exc → exc connection had a total in-degree of 200, and the exc → inh, inh → exc, and inh → inh connections had a in-degree of 400. These incoming connections were distributed homogeneously over all compartments. Finally, the Poisson input generators – with firing rates of 400 Hz – where connected in a one-to-one fashion to the somata of the excitatory neurons.

**Ca-AdEx simulations.** The Ca-Adex model featured the dynamics shown in Fig 3B, with parameters adapted from Pastorelli et al. [17]. Additionally, appropriate input receptors were added for each simulation paradigm. Specifically, for the BAC firing protocol, we had a somatic input current receptor port which received a short current pulse of 5 ms duration and 1000 pA amplitude, and a dendritic current-based synapse with a dual exponential profile (*τ_r_*= 0.2 ms, *τ_d_* = 10 ms) of weight 400 pA. To simulate the STDP protocol, we used an AMPA synapse with an initial weight of 10 nS and a maximal weight of 20 nS. Other STDP parameters can be consulted in the code (see also Fig S6).

**Winner-take-all network.** The network described as a use-case for the NESTML compartmental modelling framework represents a specific customization of the more general WTA-states model. In this model, based on a Winner-Take-All (WTA) mechanism, populations of recurrently connected excitatory neurons, the neural assemblies, are able to learn memories through the combination of a specific contextual signal with a sensory input. The WTA mechanism is obtained through competition among populations of excitatory neurons connected towards the same population of inhibitory neurons.

The model, implemented in NEST, can be configured to work with both two-compartment (Ca-AdEx) and point-like (AdEx) excitatory neuron models, showing how the performance takes advantage from the apical mechanism in the Ca-AdEx model. Several parameters of the network, such as the number of neurons in each population and the number of populations, can be defined by the user through configuration files in yaml format. In this work, the WTA-states network was configured with four excitatory populations, each one composed of 20 recurrently connected neurons. A single inhibitory population of 20 neurons is connected to all the excitatory populations, implementing the WTA-dynamics. The connectivity, both recurrent intra-population and between excitatory and inhibitory neurons, is implemented using *all-to-all* connections with the following synaptic weights: *w*_exc→exc_ = 12 nS, *w*_exc→inh_ = 7.0 nS, *w*_inh→exc_ = 1.5 nS, *w*_inh→inh_ = 1.0 nS, where the synaptic receptor currents follow the dual exponential profile (5) with *τ_r_*= 0.173 ms, *τ_d_* = 0.227 ms, and *e_r_* = 0 mV for the excitatory synapses and *τ_r_* = 1.73 ms, *τ_d_* = 2.27 ms, and *e_r_* = −85 mV for the inhibitory synapses. In addition, excitatory populations are stimulated with a Poisson noise: each neuron receives input from 20 Poisson generators firing at 600 Hz, trough a synaptic weight *w_noise_* = 1.0 nS. Note that the passive electrical parameters of the point-neuron were tuned to have the same input resistance as the two-compartment neuron. However, the two-compartment model had additional membrane leak not incorporated in this analysis because of the Ca^2+^-activated K-current, resulting in slightly lower overall network firing rates for identical weights. To correct this, we increased *w*_exc→exc_*, w*_inh→exc_ and the external input weight with a scale factor of 1.1 in the two-compartment case, which resulted in approximately equal firing rates for the corresponding populations.

New memories were written into the network using a contextual signal in the form of a Poisson stimulus of 3000 Hz during a 25 ms window, injected trough dual exponential current-based synapses with *τ_r_*= 0.173 ms, *τ_d_* = 10 ms, and synaptic weight *w_stimulus_*= 4.0 nA, into selected excitatory populations. This excitatory stimulus was preceded by 200 ms of inhibitory Poisson input at a rate of 200 Hz, through conductance-based synapses with *τ_r_* = 1.73 ms, *τ_d_* = 2.27 ms, *e_r_* = −85 mV, and weight 2 nS. Recurrent connectivity in the targetted populations was made plastic through a multiplicative STDP model [95] with *λ* = 0.1, *α* = 1.05, *τ*_+_ = 20 ms.

All connections impinge the perisomatic compartment, except for the contextual signal that, only when using two-compartments Ca-AdEx neurons, target the distal compartment allowing for the activation of the apical amplification mechanism.

Note that the contextual stimuli were identical in the two-compartment and point-neuron networks.

## Supporting information

### Code concepts

In the new compartmental modelling framework, the coding of a network simulation featuring dendritic compartments broadly speaking consists of three steps: first, the subcellular dynamics are described in NESTML (Fig S3, top), second, the models are built in the PyNEST API (Fig S3, middle), and third, the models are connected to each other, and to the necessary input generators and recording devices (Fig S3, bottom).

**Defining ion channels and receptor dynamics.** The passive system in eq. (1) is always assumed to be present, and does not need to be explicitly stipulated in NESTML. Rather, the active currents that feed into it are user-described. These currents can read out the compartmental voltage, which in NESTML is described through the `v_comp` keyword. The compartmental code generator interprets two broad categories of currents: channel currents (Fig S3, top left), which are defined as `inline`s tagged with the @mechanism::channel tag, and receptor currents (Fig S3, top right), which are `inline`s tagged with the @mechanism::receptor tag. The only semantic difference between them is that receptor currents integrate inputs defined in the `input` block. These current types are defined respectively through the `channel` and `receptor` keywords in the `equations` block in NESTML, and associated parameters and state variables are defined respectively in the `parameters` and `state` blocks. Describing the evolution equations for the state variables then follows the normal NESTML syntax [29]. We note that the integration of spiking inputs by receptors can be described through the `kernel` syntax element, as in (Fig S3, top right), or through an explicit `onReceive` statement, as in Fig S4A.

**Constructing models in the PyNest API.** The currents described in NESTML are the ones that will be available to include in the compartments of the model when it is constructed. At runtime, in the PyNEST API, the model is constructed by initializing its `compartments` attribute from a list of dictionaries, that (i) encapsulate the dendritic tree structure by stipulating a parent index for each compartment, and (ii) that feature a dictionary of parameter values describing both the parameters of the passive system (cf. eq. (1)) and the NESTML-defined channel mechanisms (Fig S3, middle left). To reference the compartments later on, each compartment is assigned an index that corresponds to its position in the compartments list. When certain parameters are not stipulated in the parameter dictionary, the default values defined in the `parameters` block of the NESTML code are used. Note that channel currents are first analysed by the code generator, and will only be simulated if the parameters are such that the current is not zero (e.g. if the maximal conductance is zero, the current will not be simulated in that compartment). In this way, one can flexibly customize the set of currents that will be simulated in each compartment.

After the compartments have been set, receptors are added to the model by setting the `receptors` attribute (Fig S3, middle right). Receptor currents differ from ion channels, in that there can be at most one ion channel of a given type per compartment, but there can be an arbitrary number of receptors. Receptors are also initialized as a list of dictionaries, each individual dictionary featuring the index of the compartment it targets, the receptor type, which correspond to one the NESTML-defined names of the receptor currents, and optionally a parameter dictionary. To connect inputs to the receptors, the standard NEST syntax is used (Fig S3, bottom left), where the receptor type corresponds to the index of the targeted receptor in the receptors list. Finally, to record state variables in the model, the standard multimeter is used, where the NESTML-defined name of each state variable is suffixed either by the compartment index – if the state variables is associated to a channel – or the receptor index – if the state variable is associated to a receptor (Fig S3, bottom right).

**Integrate-and-fire type mechanisms.** To implement integrate-and-fire type mechanisms, users can leverage the NESTML `update` and `onReceive` blocks. The update block allows for the insertion of arbitrary control statements that govern the evolution of the system, but are not part of the smoothly evolving ODEs defined in the `equations` block. This includes, for instance, `if-else` statements that measure threshold crossings. To describe dynamics that depend The `self_spikes` variable has a special meaning in NESTML, and signifies the neuron’s own spikes. We use it here in the `onReceive` block, to trigger the initiation of the absolute refractory period and the addition of the spike-triggered adaptation (Fig S4A). Accordingly, we use the update block to measure the duration of the absolute refractory period. We note that the ongoing dynamics of the model can not be paused during the absolute refractory period, as is the case with point neurons, as the dendritic compartments still integrate inputs normally. For this reason, we implement the refractory period as a channel mechanisms (`refr` in Fig S4A) that clamps the voltage to the reset value (Fig S4D, top), following the original formulation of the Ca-AdEx model that was directly written in C++ code for NEST [17]).

To simulate the AdEx dynamics only at the soma, we provide the desired parameters to the somatic compartment during model construction in the Pynest API, but set the maximal conductances to zero in the dendritic compartment (Fig S4B), resulting in a model with a passive dendritic compartment and AdEx somatic compartment (Fig S4C). Finally, we illustrate the resulting dynamics by targeting the somatic and dendritic compartment with excitatory and inhibitory Poisson inputs (Fig S4D), confirming (inset) the correct generation of somatic spikes along with the refractory period and reset, decoupled somatic and dendritic dynamics (top), the correct behaviour of the spike-triggered adaptation (middle) and of the exponential threshold current (bottom).

**Concentration dynamics.** To introduce slow concentration dynamics, equation or set of equations can be marked with the @mechanism::concentration tag. In doing so, we effectively instruct the code generator that they can be factored out, and that mechanisms reading the associated variables can effectively consider them to be constant for the duration of an integration step. Without it, the code generator would effectively have to consider the calcium current, potassium current and slow mechanism as a single system of *N*_tot_ = *N*_chan_Ca_ + *N*_chan_K_+ *N*_c_Ca_ state variables, requiring e.g. propagation matrices of size *N*^2^. Through the use of the @mechanism::concentration moniker, the code generator knows that this can be split up in three systems of respective sizes *N*_chan_Ca_, *N*_chan_K_, and *N*_c_Ca_. By extension, the @mechanism::concentration keyword allows the efficient incorporation of various processes that proceed on a slow time-scale compared to the fast channel integration, allowing for instance the simulation of various molecular signalling pathways.

**Combining plasticity processes and compartments.** To implement plastic synapses, several options exist in the NESTML compartmental modelling framework. First, it is a reasonable strategy, especially when the plasticity rule follows a complex interplay of postsynaptic factors, to implement the associated set of equations as part of the compartmental model, leveraging the features described previously. Secondly, one could also create NESTML-defined synapse models, or use existing ones, and compile them together with the compartmental model (Fig S6A, top). This results in the generation of receptor currents that feature the desired plasticity process as part of their dynamics, for every receptor current implemented in the NESTML compartmental model. The name of this receptor current then follows the convention `{receptor name}_{plastic synapse name}_nestml` (Fig S6A, bottom left).

When following these two strategies, the connection should be initialized as a static synapse in the PyNEST API, as the plasticity dynamics are simulated entirely on the post-synaptic side. Finally, if the plasticity model is part of the standard model collection in NEST, it can also be specified when the connection is initialized (Fig S6A, bottom right). In this case, the receptor itself should be the static version.

## Supplementary figures

**Figure S1.**
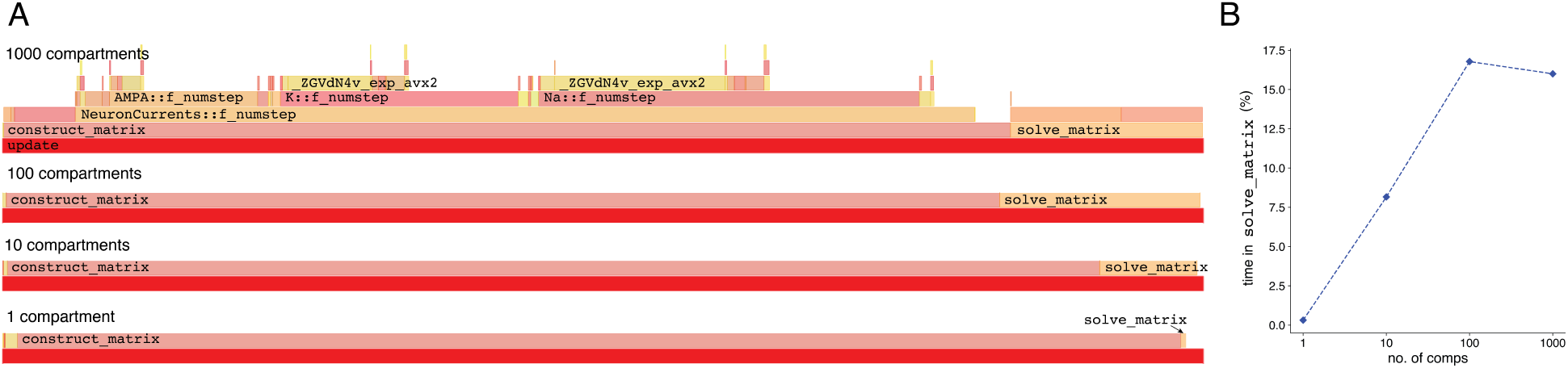
Profiling results for the MC framework. **A:** Profiling graphs of the runtime correlate extracted with the Hotspot profiler^2^, quantifying the relative fraction of time spent in specific function calls in the linear dendrite model from Fig 2A. We highlight the `update` function (red), which implements the full update step of the model. This function is broken down in a `construct_matrix` part and a `solve_matrix` part, which respectively construct and solve the Hines matrix. Through the `construct_matrix` function all relevant update functions are called. Here, there are an AMPA receptor, a K channel and a Na channel in each compartment (shown for the 1000 compartment case, top). Decomposing these updates reveals that (i) these functions are vectorized properly as the vector primitive of the exponential function is invoked and (ii) that they spend most of their time evaluating exponentials to compute the channel activations and advance the channel state variables. Profiling results were obtained an a laptop with Intel(R) Core(TM) i5-4210H CPU @ 2.90GHz and 16 GB DDR3 1600MHz Ram **B:** Quantification of the fraction of time spent solving the Hines matrix (i.e. in the `solve_matrix` function) vs. the combined time spent to construct and solve the matrix (i.e. `construct_matrix` + `solve_matrix`) as a function of the number of compartments.

**Figure S2.**
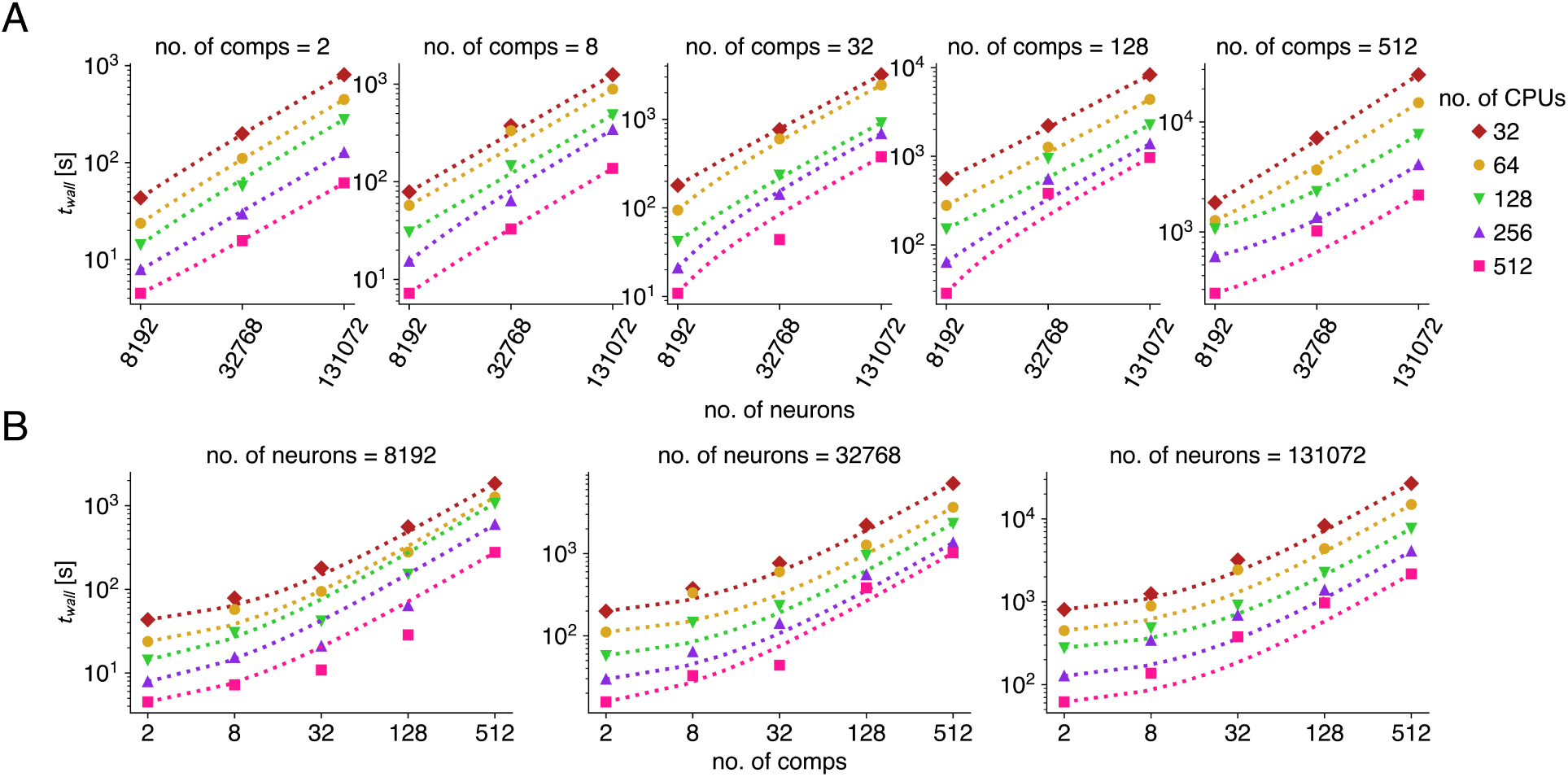
Runtimes of the network scaling benchmark. **A:** Runtimes increase linearly as a function of the number of neurons in the network. **B:** Runtimes also increase linearly as a function of the number of compartments in the network. Note that this does not result in linear plots on a loglog scale as the *y*-value of the zerocrossing of the linear relationship (no. of comps = 0) is not at zero. The dotted lines show linear interpolations between first and last points.

**Figure S3.**
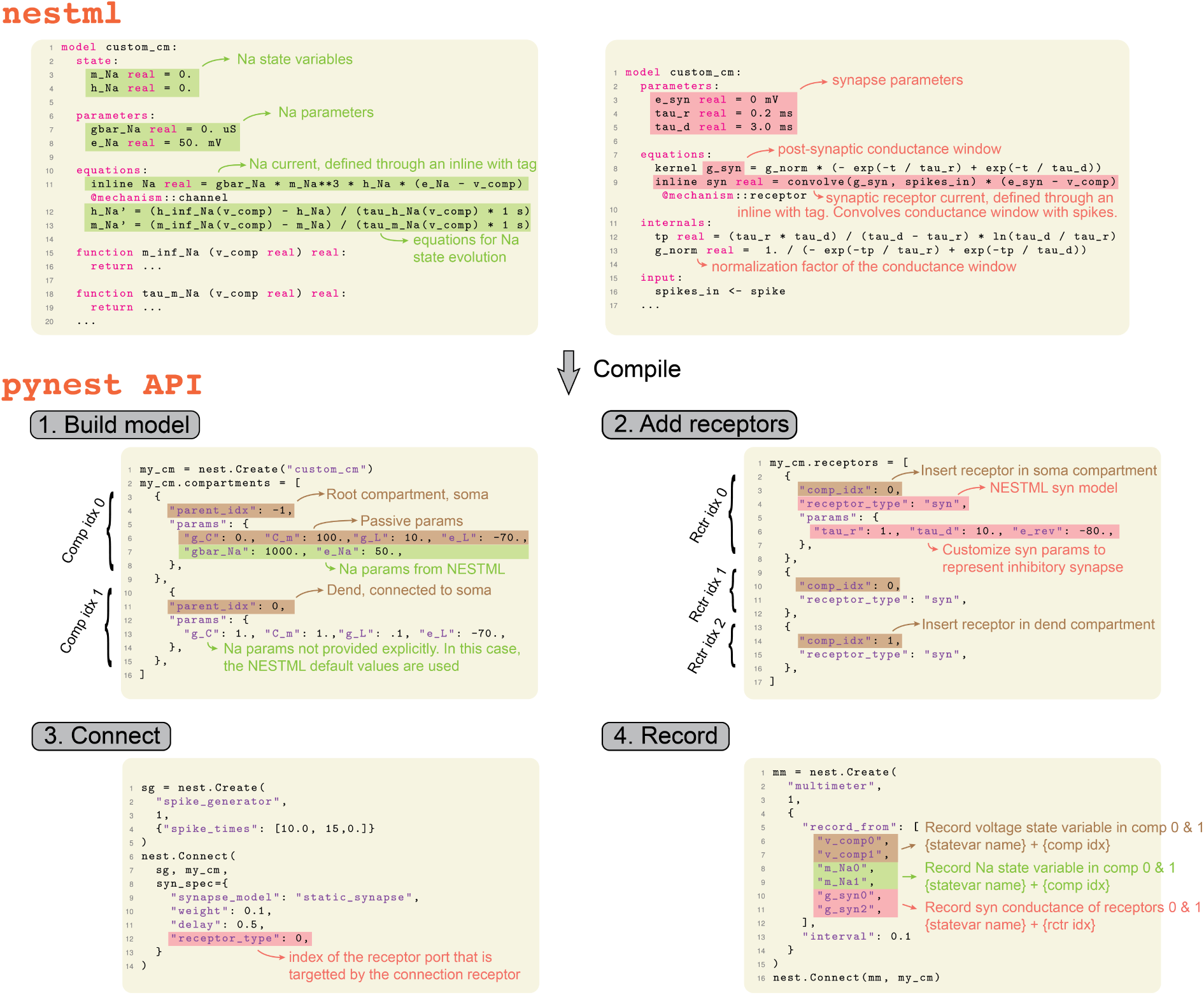
Basic code concepts in NESTML and the PyNEST API. Ion channel currents (upper left) are defined through `inline`s tagged with the @mechanism::channel tag. The current consist of state variables and parameters, which are defined in the respective NESTML blocks. State variable evolution equations follow the standard NESTML syntax. Receptor currents (upper right) are defined in a similar manner as channels, but are tagged with the @mechanism::receptor tag, and should integrate some input (here a spiking input, but continuous inputs can also be integrated – see documentation). To build the model in the PyNEST API (middle left), an empty model is initialized with `nest.Create()` and then equipped with compartments. For each compartment, custom parameter values or state variable initializations can be specified in the `“params”` dictionary. The compartments get assigned an index based on their position in the compartments list (‘Comp idx’ in the figure). Note that the parent compartment (specified with `“parent_idx”`) can only refer to a compartment at a prior position in the `compartments` list, and that the somatic compartment (the ‘root’ node of the tree) should have `-1` as parent index. After the compartments have been set, the receptors can be added (middle right) by specifying the compartment index that they target, the type of receptor (one of the types defined in NESTML) and the parameters and state variable initializations. Similarly to compartments, receptors get assigned an index corresponding to their position in the `receptors` list (‘Rctr idx’ in the figure). Finally, the models can be connected (bottom left), specifying the index of the receptor port that a connections targets as part of the `syn_spec` dictionary (the `“receptor_type”` entry), and recordables can be set by connecting a `“multimeter”` to the neuron model (bottom right). Note that any NESTML defined state variables, inlines or kernel convolutions can be recorded.

**Figure S4.**
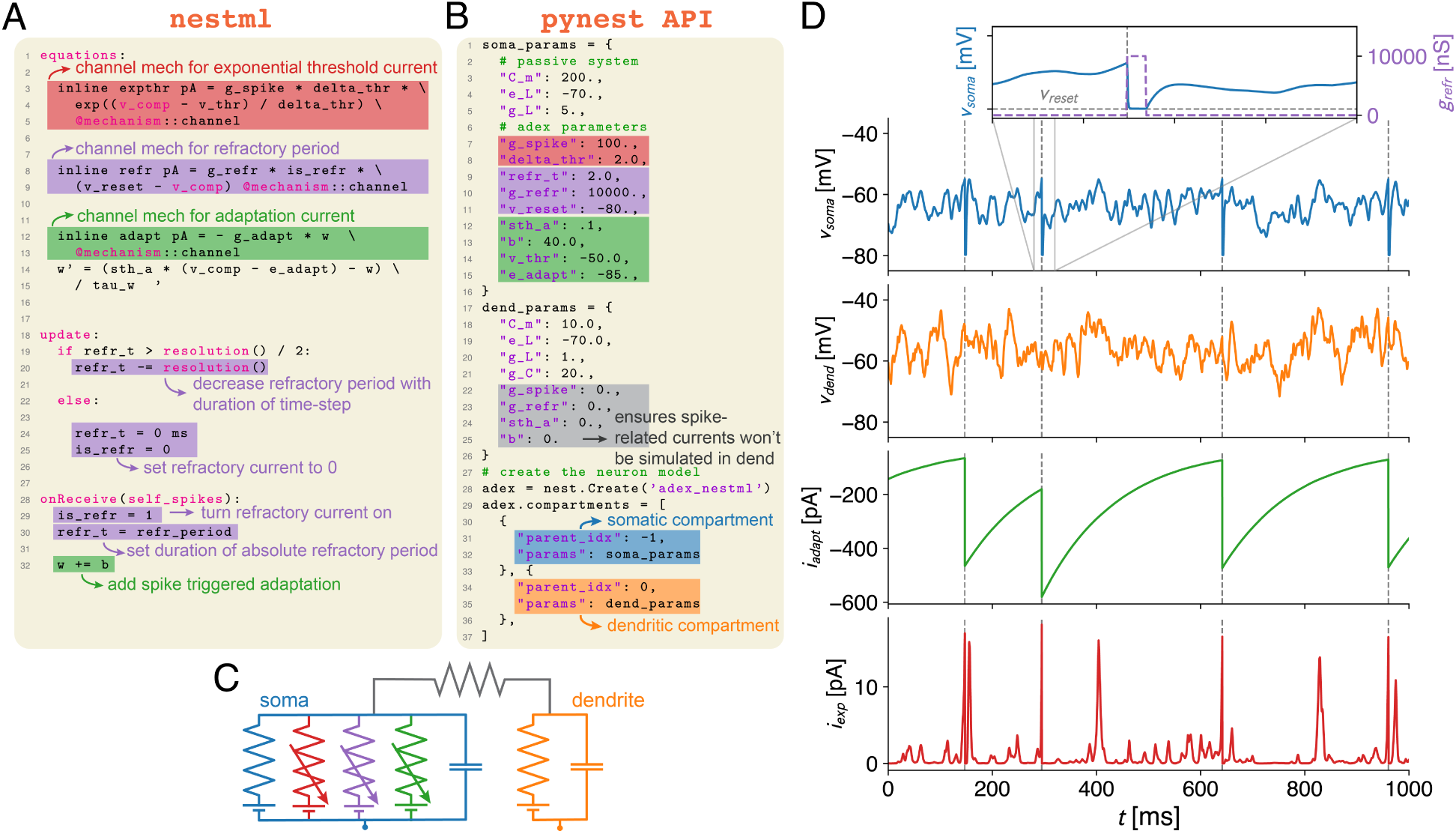
Implementing adaptive exponential integrate-and-fire (AdEx) dynamics in the compartmental modelling framework. **A:** The NESTML language allows for a compact description of non-linear trans-membrane current dynamics, here illustrated with the AdEx model. This model features an exponential threshold current (`expthr`, red) and an adaptation current (`adapt`, green). While the spike threshold is set in the PyNest API (B), spike adaptation (green), refractory period and reset (purple) dynamics are implemented on the NESTML side using the `onReceive` and `update` blocks. As compartmental dynamics are always ongoing, the absolute refractory period is implemented by clamping the voltage to the reset potential. Note that these dynamics defined in NESTML represent transmembrane currents that *can* be implemented in each compartment of the eventual model, but whether they *will* be simulated in the compartment depends on the final parameter choice. **B:** At model creation in the PyNest API, compartment parameters are set to their final values (note that parameters not explicitly defined in the `“params”` dictionaries are initialized to their NESTML default values). Here, in the dendritic compartment (orange), conductance parameters for the AdEx currents (`expthr`, `adapt`, and `refr`) are set to zero, while in the somatic compartment (blue) they are set to non-zero values. **C:** Schematic of the resulting model. Note that because the conductance parameters of the AdEx mechanisms were chosen such that these currents are identically zero in the dendritic compartment, they are *not* simulated there. **D:** Exemplar simulation of the resulting model, stimulated with somatic and dendritic Poisson inputs, illustrating (from top to bottom) the somatic voltage with an inset detailing the refractory period implementation, the dendritic voltage, the `adapt` current and the `expthr` current.

**Figure S5.**
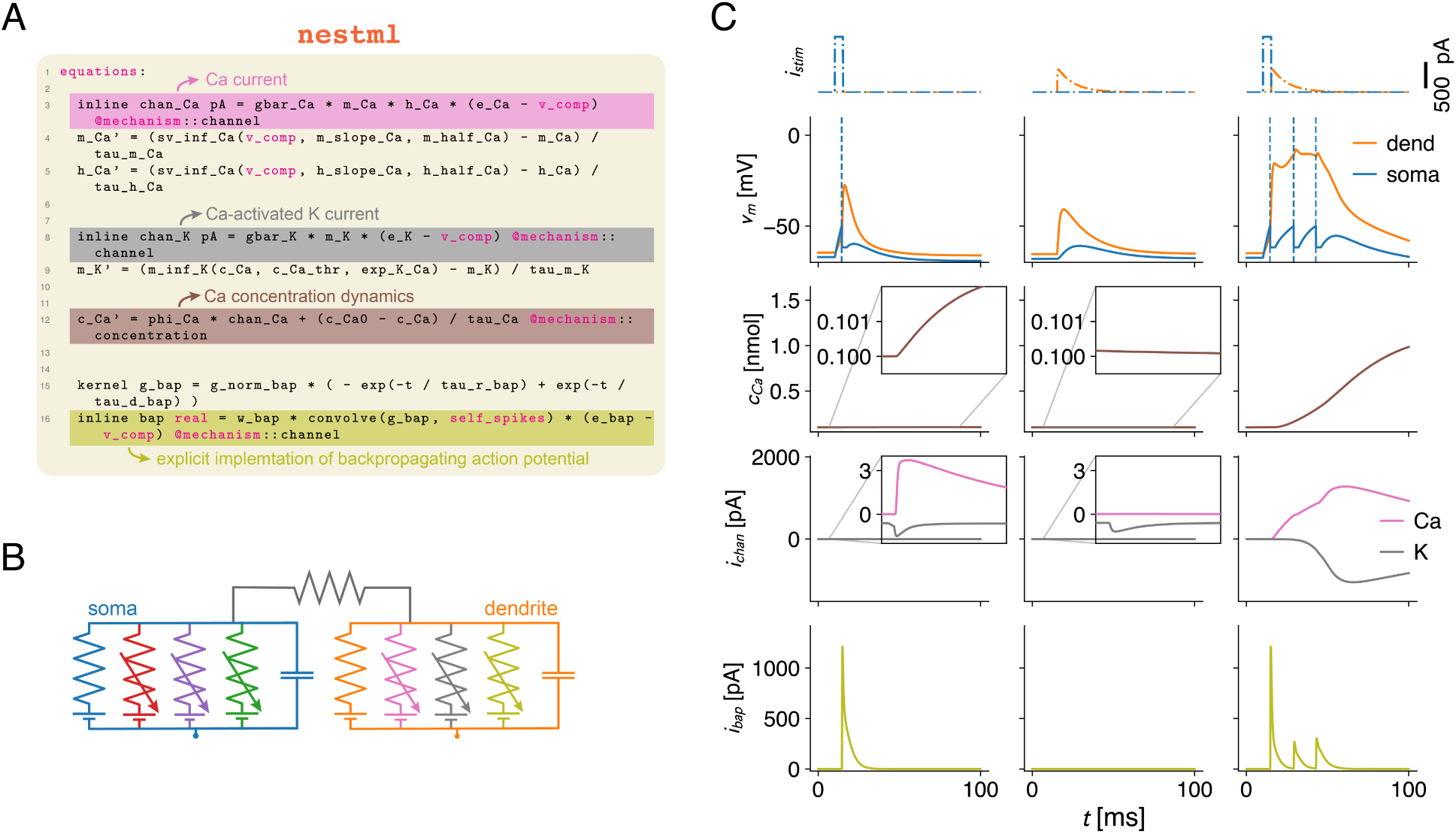
Combining somatic AdEx and dendritic Ca^2+^ dynamics. **A:** In the NESTML framework, transmembrane currents (here a Ca^2+^ and a K^+^ current, resp. pink and grey) can interact with slow mechanisms, such as e.g. a mechanism modelling the Ca^2+^ concentration, using the @mechanism::concentration keyword (brown). Here, we also add an explicit implementation of the bAP (`bap`, yellowgreen), leveraging the `self_spikes` keyword. **B:** Schematic of the final model, featuring the AdEx currents in the somatic compartment, and the Ca^2+^, K^+^, and bAP currents in the dendritic compartment. **C:** Simulation of backpropagation-activated Ca^2+^-spike (BAC firing) in the resulting model, where coincidence of a somatic input eliciting a single AP in isolation and a dendritic PSC waveform (top) results in the generation of a dendritic Ca^2+^-spike and somatic AP burst (second from top). Subsequently, the increase in Ca^2+^ concentration (middle) through the large Ca^2+^ current activates the K^+^ channel (second from bottom), repolarizing the dendritic compartment. To obtain this interaction, the bAP current is implemented explicitly (bottom) as this simplified model lacks Na^+^ channels along the apical dendrite to boost the bAP.

**Figure S6.**
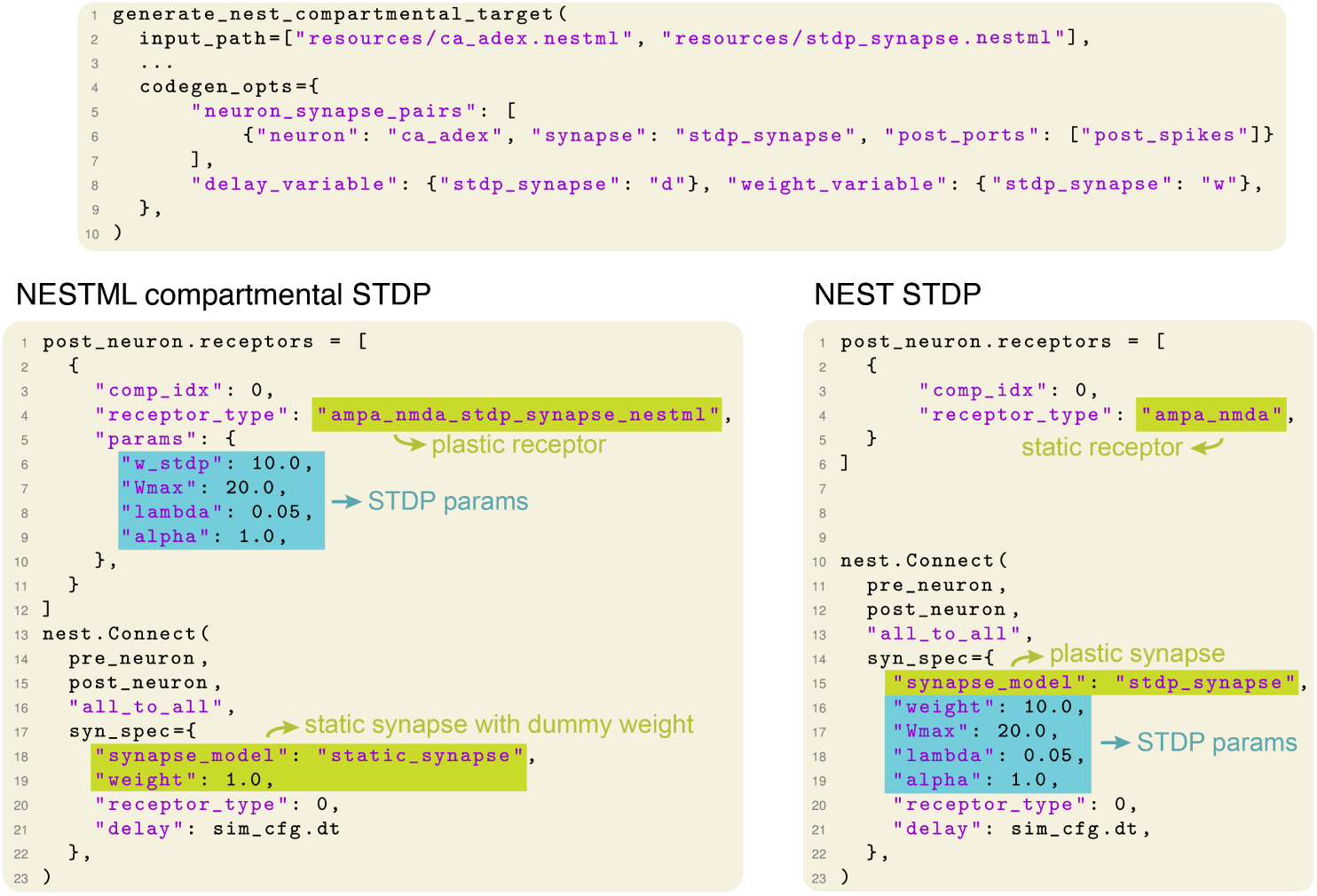
Adding plasticity synapses to the multi-compartment models. NESTML-defined plasticity rules can be compiled together with a neuron model (top). In this case, the plasticity rule is combined with every receptor current defined in the neuron model. Note that the pure receptor current – without plasticity – is also still available. In the PyNEST API, this plastic receptor is added when initializing the `receptors` list, whereas the connection itself is initialized as being static (bottom left). Standard plasticity rules already defined in NEST can also be used, in this case, the original receptor is added in the `receptors` list, but the connection is initialized as a plastic synapse (bottom right).

